# Functional maturation in abducens motoneurons populations during angular VOR larval development

**DOI:** 10.64898/2026.07.28.741290

**Authors:** Mathilde Pain, Marie Boulain, Laura Cardoit, Marie-Jeanne Cabirol, Jean Forgue, Martin Gorges, Gilles Courtand, Stefan Glasauer, François M. Lambert

## Abstract

Extraocular motoneurons are the final neuronal relay implicated in gaze motor control and are known to be subdivided in functional subgroups, differently implicated in ocular motion dynamics. However, the maturation of these functional populations of extraocular motoneurons, in relation with the development of gaze-stabilizing reflexes remains largely unexplored. In amphibian tadpoles, the angular vestibulo-ocular reflex (VOR) appears later than other visuo-vestibular ocular reflexes and matures until the metamorphosis climax. Two types of Abducens motoneurons have been described to participate to the angular VOR in larval frog: spontaneous motor units, exhibiting a robust resting activity and silent motor units recruited only during head motion. The aim of this study was to investigate the maturation of these two types of Abducens motor units in relation with the development of the angular VOR by evaluating their discharge dynamic in response to head rotation in semi-intact preparations of larval *Xenopus laevis*. During larval life, the discharge modulation during sinusoidal head rotations increases significantly for silent units only, demonstrating a better sensitivity of this Abducens motoneuron sub-population to horizontal semicircular canal activation. In addition, this functional maturation was accompanied by an increase of the myelination in the lateral rectus motor nerve, promoting a faster conductivity in late larval stages than in early one. These findings showed that the development of the angular VOR is supported by a selective maturation of extraocular motoneurons subpopulations, specifically implicated in the improvement of the ocular kinematic during the reflex.

## INTRODUCTION

Gaze stabilization results from synergistic interactions of sensory feedbacks and internal feedforwards inputs within dedicated circuits ensuring appropriate transformation of multi-modal signals into a single motor output for compensatory eye movements. Classically, optokinetic and vestibulo-ocular reflexes (OKR, VOR) guarantee this transformation during passive head/body motion (Angelaki et Cullen 2008) and are supplemented by locomotor-induced gaze responses (LGR) produced by CPG efference copy commands during locomotion (Lambert et al. 2012). Extraocular motoneurons constitute the final neuronal output for all of these sensory-motor mechanisms and convey the appropriate motor commands responsible for the ocular response. These motoneurons appear to form distinct functional subpopulations, that are more or less involved in the large diversity of ocular movements (Delgado-Garcia et al., 1986.; Dieringer et Precht 1986; Mays et al. 1991; Gamlin et Mays 1992; Davis-López De Carrizosa et al. 2011; Carrero-Rojas et al. 2021; Dowell et al. 2025; for review see Horn et Straka 2021). A previous study in larval frog revealed that two subtypes of abducens motoneurons, distinctly separated by their firing activity at rest, participate conjointly to the angular VOR: spontaneously active and silent motor units, respectively (Dietrich et al. 2017). These two subtypes of abducens motor units were described to encode differently head rotation signals in the horizontal plane.

During the ontogeny, sensory-based and intrinsic gaze-stabilizing ocular behaviors do not develop through the same temporal pattern, depending on species specificity and eco-physiological constraints (Faulstich et al. 2004; Beraneck et al. 2014). Larval amphibians and fishes acquire OKR, otolith VOR and LGR early during the development, few days after hatching (Beck et al. 2004; Lambert et al. 2008; Bianco et al. 2012; Schneider-Soupiadis et al. 2025; Leary et al. 2025). Inversely, the duct size limitation imposed a low sensitivity for semicircular-canals and constrains the angular VOR being functional later during the development (Beck et al. 2004; Lambert et al. 2008; Schneider-Soupiadis et al. 2025). Following its developmental onset, the canal-driven VOR seems to undergo a second maturation stage, that was simply reported by a significant increase of the reflex gain so far (Bacqué-Cazenave et al. 2022; Schneider-Soupiadis et al. 2025). However, the precise consequences of this VOR maturation on the ocular dynamics in one hand, and concomitantly on the two populations of abducens motor units in another hand, remains to be determined. More importantly, investigating the developmental plasticity of these two abducens motor unit subtypes during the VOR maturation is important for a better understanding of how distinct ocular kinematics are produced by separate extraocular motor neurons subsets, as recently explored in larval fish (Dowell et al. 2025).

Here, eye movement dynamic and abducens motor nerve discharge elicited by natural activation of horizontal semi-circular canals were measured during Xenopus larval life, from early stage with a weak angular VOR, to older one, just before the climax of the metamorphosis. Our study demonstrated that the VOR maturation relied on a better control of the eye velocity supported by an improvement of the discharge dynamic of the silent motor units, the subtype of abducens motoneurons activated during head movements only and encoding specifically for the velocity parameter. Reversely, the response of spontaneous motor units remained unchanged. Such functional changes were also associated to morphological modifications of myelinated axons in the abducens nerve and a better innervation of the lateral rectus muscle, contributing to a faster and a broader activation of the extraocular muscle fibers.

## MATERIAL AND METHODS

### Animals

Experiments were performed on *Xenopus laevis* larvae, at developmental stages 49-50, 52-53 and 55-57. Animal stage was identified from morphological external criteria characterized by Nieuwkoop and Faber (1956). The animals were maintained in filtered water at the INCIA laboratory facility (EU0568; C33-063-923) in a 12-hour light/dark cycle at 20-22°C. All procedures were approved by the local ethics committee (protocols #2016011518042273 APAFIS #3612), in accordance with European regulation.

### Semi-intact preparations and semicircular canal measurements

Larvae were anesthetized in a 0.05% tricaine methanesulfonate solution (MS-222, Sigma) and secured on a sylgard bottom of a petri dish with insect needles (Entomoravia - Austerlitz Insect pins; 0.2mm), filled with oxygenated cold Ringer’s solution (93.5 mM NaCl, 3 mM KCl, 30 mM NaHCO3, 0.5 mM NaH2PO4, 2.6 mM CaCl2, 1 mM MgCl2 and 11 mM glucose, pH 7.4). The viscera and the skin covering the head were removed to expose the otic capsules and the brain. The optic nerve was cut bilaterally to eliminate visual inputs. Brightfield images of the horizontal semicircular canals were acquired from the top with an Infinity 3 camera (Lumenera) mounted on macroscope SMZ18 (Nikon). Canal size dimensions were measured using the freeware FIJI (https://fiji.sc/). Landmarks were placed manually to measure the small and the large circuit radii and the lumen radius as extensively described in (Lambert et al. 2008; Schneider-Soupiadis et al. 2025).

### Measurement of the vestibulo-ocular reflex

The petri dish containing the larval semi-intact preparation was positioned at the center of a motorized vestibular apparatus (Technoshop COH@BIT, IUT de Bordeaux, Université de Bordeaux). The preparation was constantly perfused in oxygenated Ringer solution (same than above enriched with 11.15 mM CaCl₂, 3.05 mM MgCl₂; pH 7.4) at 18°C. Horizontal rotations at different motion velocity patterns (sine or triangle step) provided natural stimulations of intact semicircular canals. Bilateral eye movements were recorded online using a high-speed camera at 100fps (Basler AG, ac1920) positioned above the semi-intact preparation. Using the same methodology than in (Bacqué-Cazenave et al. 2022), the VOR was recorded online with a homemade Python-based software (v3.8.13; https://github.com/gillescourtand/xenopusProject). Eye position was analyzed off-line with Dataview (https://www.st-andrews.ac.uk/~wjh/dataview/) to obtain the eye velocity and calculate the VOR dynamic parameters (eye velocity gain, phase relationship). In a second set of experiment, galvanic vestibular stimulation (GVS) was applied to evoke VOR without endolymph displacement in the canal. Silver wire electrodes were positioned at the surface of the otic capsule, the closest of the horizontal canal cupula the same way as previously described in (Gensberger et al. 2016). Sinusoidal current trains (200-300 µA; 0.5-1.0 Hz) were applied to activate horizontal canal nerve afferences and allowed to record a GVS-evoked VOR.

### Retrograde labeling of abducens motoneurons and BrdU pulse chase

Tadpoles were incubated at stage 49-50 in water with 5 mM BrdU (Sigma) for 24 hours in the dark (PULSE). After the BrdU pulse, animals were rinsed three times to remove completely BrdU traces and were bred until stage 55-57 (CHASE) where abducens motoneurons were retrogradely labeled. abducens motoneurons were labeled in semi-intact preparation by application of retrograde dextran amine tracers in the lateral rectus muscle (Life Technologies; 568nm). 3-5 dye crystals were applied in a small cut done at the muscle nerve junction, like in Dietrich et al (2017). The excess of dye was washed out with Ringer’s solution then the preparation was incubated in oxygenated Ringer solution overnight at 14-16°C to allow the retrograde tracer migration into the motoneuron soma before histological treatments.

### Immunohistochemistry

Consecutive to the fluorescent tracer migration, the hindbrain was dissected out from the preparation and fixed overnight at 4°C in 4% paraformaldehyde (PFA, Sigma-Aldrich) made in a 0.1% phosphate buffer saline solution (PBS). After fixation, tissue was incubated in a 20% sucrose PBS solution for 24 h. Samples were embedded in Tissue-Tek (VWR Chemicals) and frozen in isopentane (Avantor VWR) at −45°C. Frozen hindbrains were sliced at -20°C with a cryostat (CM3050, Leica) to obtain series of 25μm cross-sections. As a prerequisite for BrdU immunodetection, slices were incubated in a 8% HCl solution, followed by 0.1 M borate buffer to neutralize the acid, facilitating the access of the BrdU antibody to the nucleus. Following the saturation step with PBS, 10% Donkey serum (Abcam), and 0,3% Triton X-100. Cross-sections were incubated overnight with a rat primary anti-BrdU (abcam) at 1:500 and revealed with a donkey secondary anti-rat at 1:500 for 2 hours. Finally, cross-sections were mounted with Dapi fluromont-G (Southern Biotech).

Lateral rectus muscles together with the connected abducens nerve branch were dissected out from stages 49-50, 52-53 and 55-57 (see preparation procedure above) and fixed in a stretched position during 2 hours in 4% PFA at room temperature. Samples were rinsed three times in PBS and incubated 1h30 at RT in a blocking solution containing PBS, 1% bovine serum albumin (BSA) (Sigma), and 0,3% Triton X-100. Muscle tissues were incubated overnight at 4°C with a mouse anti-3A10 (1:300, supernatant; DSHB), labeling neurofilament and with the Alexa Fluor-conjugated α-bungarotoxin (1:1500, Alomone), labeling postsynaptic acetylcholine receptors at the neuromuscular junction. Samples were rinsed the next day in PBS and incubated with species-specific fluorescent secondary antibodies (1:500, Invitrogen) for 2 hours at room temperature. Lateral rectus muscles histological preparations were mounted in concave slide with fluromont-G (Southern Biotech).

### Confocal fluorescent image acquisition and analysis

Fluorescent images were acquired with a LSM900 confocal laser microscope (Zeiss) with 488, 543, and 633 nm laser lines. Multi-image Z-stacks of abducens motoneurons were acquired with a 20×/0.8 objective at a 1.5μm image interval. Lateral rectus muscles images were acquired with a 20×/0.8 or a 63x/1.4 for detailed views with a 1.5μm image interval. Neuronal population or muscles images were obtained through orthogonal projections of confocal image stacks. The final images were artificially colored and processed in FIJI and Adobe Photoshop (Adobe Systems, Inc., San Jose, CA, USA). For each animal, one lateral rectus muscle was randomly selected, and the entire muscle volume was imaged. Image stacks were processed using a Top Hat filter followed by a threshold adjustment. Contrast was normalized for each slice. The region of contact between the nerve and the muscle was identified by an α-bungarotoxin positive signal corresponding to the synaptic labelling. Three-dimensional volume measurements of this region were performed using the 3D ImageJ Suite plugin (doi:10.1093/bioinformatics/btt276).

### Lateral rectus motor nerve electrophysiological recordings

Semi-intact preparations were performed for electrophysiological recording the same way than described above. The abducens nerve branch innervating the lateral rectus muscle was cut at the entrance in the muscle and suctioned in a borosilicate glass electrode (tip diameter ∼100 μm; GC120F, Harvard Apparatus) filled with Ringer’s solution. The discharge of the abducens nerve branch was recorded in response to either one-way step or sinusoidal head motion in oxygenated Ringer solution at 18°C. Nerve signal was amplified with an EXT 10-2F amplifier (NPI Electronic) and digitized with a CED Micro1401 interface (Cambridge Electronic Design) and the Spike2v7 software. The multi-unit recordings of the abducens motor nerve discharge were analyzed offline with Spike2 and Dataview.

### Data analysis of multi-unit and single-unit activity

Amplitude thresholds were set to separate motor units with a spontaneous activity from motor units that were silent or barely discharging at rest. From this thresholding, spike events were separated into two channels, corresponding to the discharge of spontaneous and silent motor units, respectively. During one-way horizontal step of the head, the instantaneous firing rate (in spike/s) was calculated for each event channel. For both spontaneous and silent multi-unit discharge component, the peak response time was the time from the maximum instantaneous firing rate (the peak response) and the end of the step, the half response decrease latency was the time when the firing rate went back to half of the maximum rate after the peak. During sinusoidal head rotations, the averaged firing rate was determined with a peristimulus time histograms (PSTH) timed on the vestibular stimulus cycle and calculated over 10 cycles at least. The discharge modulation was calculated from the difference between the firing rate at the peak response during the ON phase and the minimum firing rate during the OFF phase. Single-unit analysis was performed through the spike2 and Dataview spike sorting toolbox to extract individual single motor unit activity from the multi-unit recording. The motor unit detection was based on both the shape and the amplitude of the spike and putative single units were classified with a principal component analysis (PCA). Each PCA cluster was evaluated for single-unit validity using inter-spike interval (ISI) histograms. Only units presenting separated clusters with a clear refractory period (i.e., absence of spikes within a 1-2ms window) were analyzed with the same PSTH method than the multi-unit recording. The discharge modulation was also calculated for each identified single motor unit.

### Electron microscopy

Samples preparation and imaging were performed at the electronic Bordeaux Imaging Center (https://www.bic.u-bordeaux.fr/). abducens motor nerves were fixed by immersion in 0.1 M cacodylate buffer (pH 7.4) containing 1.6-2% glutaraldehyde and 0.02% CaCl₂ for 2 hours at room temperature. Sucrose (70 mM) was added, and samples were stored at 4 °C overnight for complete fixation. Because of the small size of the samples, several drops of a 1% osmium tetroxide solution in sodium cacodylate 0.1M were added in the tubes, allowing tissues to blacken. Then, they were rinsed 3 times with 0.1M cacodylate buffer. Thereafter, samples were immersed in 1% osmium tetroxide diluted in cacodylate buffer (without CaCl₂) at room temperature for a duration of one hour. Two additional rinses with 0.1M cacodylate buffer were performed (30 minutes each) with the final rinse being performed in distilled water. Samples were dehydrated through graded acetone baths (50%, 70%, 95%, and three times in 100%, 10min each). Resin infiltration was initiated during 2 hours in a 1:1 mixture of acetone and freshly prepared epoxy resin (Epon 812). Samples were incubated overnight in pure resin without catalyst, then 2 hours in resin containing BDMA at room temperature. Resin polymerization was achieved at 60°C for 48 hours. Transversal semithin (1µm) and ultrathin (75nm) sections of the abducens nerves were realized with an ultramicrotome (Leica UC7) and collected on squared copper grids. High-magnification images of abducens motor axons were obtained with a transmission electron microscope (Hitachi H7650). The segmentation of individual axons was conducted manually in FIJI by establishing regions of interest (ROIs) on maximum intensity projections of image stacks. For each axon, surface area and perimeter were measured. To analyze myelin coverage, the fluorescence channel corresponding to myelin labeling was threshold to isolate the signal. The resulting binary masks were used to outline myelin profiles and quantify both their surface area and perimeter. This allowed comparison between the axonal and myelin contours to assess myelination levels.

### Statistics

Data are expressed as mean with SD unless stated otherwise. All statistical analyses were performed in GraphPad Prism. Normality of each data set was tested with the Shapiro Wilk test. When normality was obtained, parametric Student’s *t*-test was used to compare two groups and one-way ANOVA to compare more than two groups, followed by Tukey’s test for multiple comparisons. When normality was not obtained, the non-parametric Mann-Whitney test was used to compare two groups and the Kruskal Wallis test for more than two groups. A difference was considered statistically significant if the *p*-value was less than 0.05. All animal or sample numbers, statistical tests and p-values are indicated either in the figures or in the figure legends. The stability of the eye motion during horizontal sinusoidal rotations was evaluated by applying a 6^th^ order polynomial fit on the eye position trace at each rotation cycle during 20 cycles trials. A regression R^2^ coefficient was obtained for each cycle and was averaged for each sequence and for each animal. Significative differences of grand mean R^2^ between stage 49-50 and 55-58 groups was tested with a Mann-Whitney U test. Head *vs* eye velocity phase relationship were plotted using polar data representations in the JASP (R) freeware environment (https://jasp-stats.org/). The uniformity of the phase distributions was assessed using Rayleigh’s test, and differences in mean phase angles between stage 50 and 55 were tested using the Watson-Williams test.

## RESULTS

### Developmental maturation of the angular vestibulo-ocular reflex during larval life

Previous studies have partially documented the progressive maturation of the angular VOR in larval Xenopus (Lambert et al. 2008; Bacqué-Cazenave et al. 2022). However, the dynamics of this maturation are still poorly quantified, limited to basic measurements of the eye position during sinusoidal head rotations. Nonetheless, both eye position and velocity signals are known to be distinctly treated by VOR sensory-motor circuits, including at motor plant (Angelaki et Cullen 2008; Straka et al. 2009; Dietrich et al. 2017). Consequently, we wanted to quantify the development of the reflex according to ocular motor parameters allowing the identification of specific abducens motor unit subpopulations.. First, eye movements were recorded in semi-intact larval preparations in response to one-way step head horizontal rotations with a linear variation of the head velocity (Fig. 1A_1_). Such a stimulus induced a rapid ocular deflection on the opposite direction followed by a slower drift back to the original position (Fig. 1A_1_). This VOR-induced ocular response was much higher at stage 55-57 than stage 49-50 (Fig. 1A_2_). At constant head velocity (±40°/s), old larvae demonstrated a significantly better velocity gain than early animals regardless the rotation frequency (Fig. 1B) with a maximum gain of 0.45 ± 0.05 (mean **±**SD, *N=7*) for stage 55-57 against 0.21 ± 0.05 for stage 49-50 (*N=6*) at 1Hz (Mann-Whitney; *p* =0.0082). However, the velocity ratio between left and right eye was similar at both developmental stages (Fig. 1C; Stage 50: R^2^=0.7978; Stage: 55 R^2^=0.8387), indicating that the angular VOR produced conjugate ocular movements already well coordinated from early stage onward. VOR-induced eye movements were also evoked in response to sinusoidal head rotations to characterize the eye motion dynamics (Fig. 1D-I). Like for one-way steps, tadpoles at stages 55-57 demonstrated a better VOR-induced ocular response than at stages 49-50 during sinusoidal head rotations (Fig. 1D). Eye velocity was significantly higher in older stages than in younger one at constant velocity (60°/s) regardless the rotation frequency. The greatest difference was observed at 1Hz where stage 55-57 performed a VOR with an eye velocity of 10.32 **±** 1.193 against only 5.467 **±** 1.086 for stage 49-50 (Fig. 1E; Mann-Whitney, p=0.0082). As for the step stimulus, left and right eye moved with the same velocity during sinusoidal head rotations at both stage (St50 R^2^=0.7540; St55 R^2^=0.5592; Fig. 1.F), confirming a good bilateral ocular coordination already at the onset of the VOR. In addition to the velocity gain, the regularity and reproducibility of the ocular signal was tested by applying a polynomial fit to the eye position trace at each sinusoidal head rotation cycle (Fig. 1G_1_). VOR-induced eye movements appeared to be smoother and more reproductible through cycle repetition, with less variations of the ocular position, at stage 55-57 than at stage 49-50 as illustrated at 1Hz in figure 1G_2_. The polynomial R^2^, quantifying the quality of the ocular position fit, was significantly higher in old larvae than in early one at 0.5 and 1Hz head rotations (Fig. 1H; At 0.5 Hz, stage 49-50 mean R^2^ = 0.8 ± 0.04 at stage 49-50 (N =8); stage 55-57 mean R^2^ = 0.92 ± 0.02 at stage 55-57 (N = 8) *p* = 0.0207; At 1 Hz, stage 49-50 mean R^2^ = 0.59 ± 0.05; stage 55-57 mean R^2^ = 0.77 ± 0.08; *p* =0.0401; Mann-Whitney). This demonstrated an increase of the VOR dynamic during the larval development at least for higher frequency stimulations. This last observation was confirmed by the noticeable improvement of the VOR phase relationship from stage 49-50 to stage 55-57, with a reduced ocular shift in old larvae for 0.5 and 1Hz (Fig. 1I; At 0.5 Hz, st49-50 phase = 196.4° ± 0.348, st55-57 phase = 165.5° ± 0.358, p=0.004; at 1 Hz, st49-50 phase = 252.6° ± 0.352, st55-57 phase = 221.4° ± 0.248 p= 0.002; Watson-Williams). Altogether, these results demonstrated a substantial maturation of the angular VOR from stage 49-50 to stage 55-57. This VOR improvement was not only supported by eye movements with larger amplitude but also with a better ocular motion dynamic. As a result, the activation of horizontal semi-circular canals at older stage produced a faster and more robust compensation of the head rotation through repetitive stimulations, particularly at higher frequencies. This maturation of the VOR may indicate an increase in vestibular sensitivity and/or a maturation at the central level.

**Figure 1.**
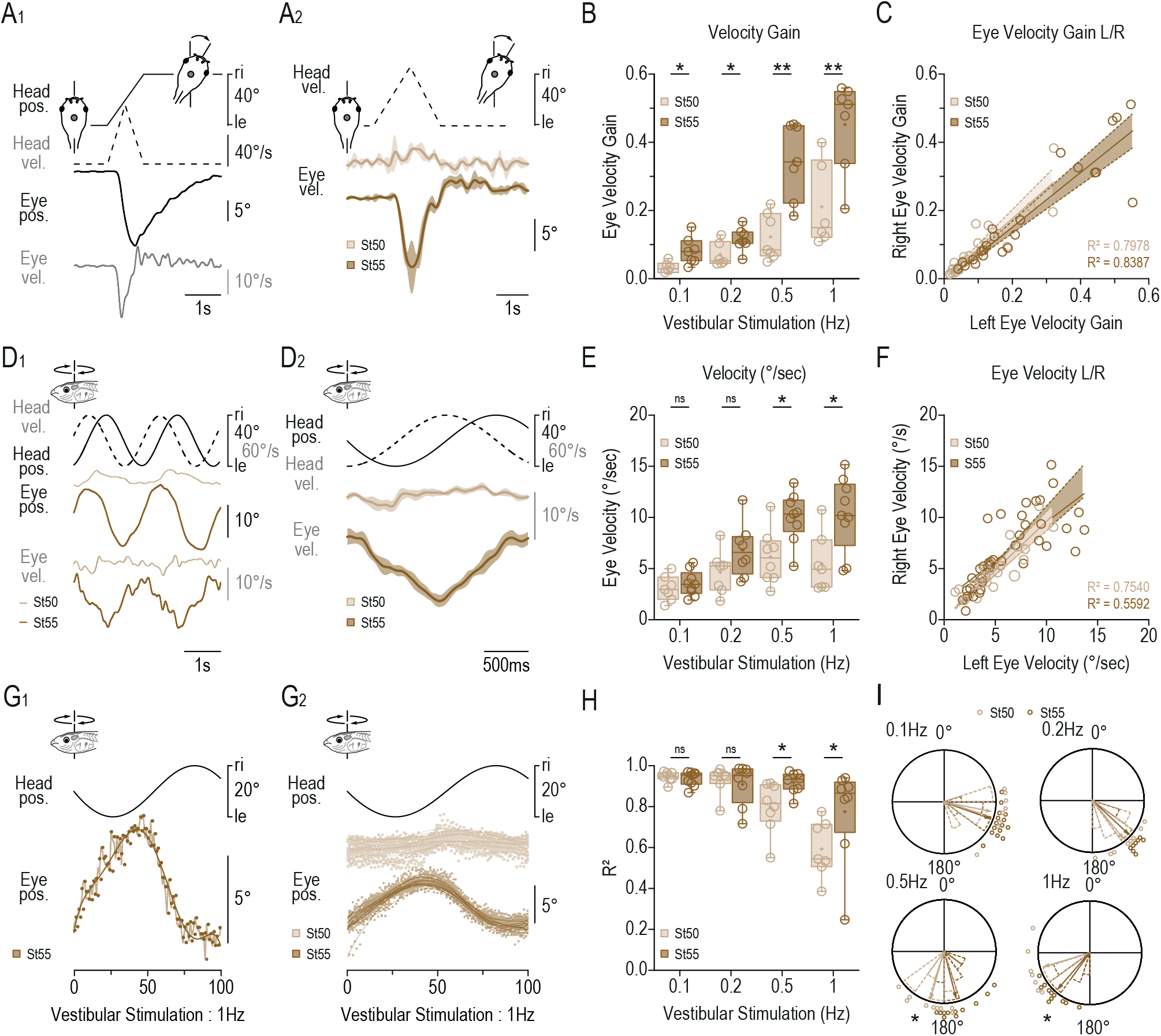
Maturation of the angular vestibulo-ocular reflex during larval life. A_1_. Example of VOR-driven eye movement in response to a rightward one-way step head positional (eye pos.) rotation at 40°, 1Hz. The eye velocity (grey, eye vel.) trace is derivate from the video-tracked eye position trace (black, eye pos.). **A_2_.** Example of averaged (±SD; 5 steps averaged) eye velocity from a stage 49-50 and a stage 55-57 larvae in response to a rightward one-way step head (eye velocity) rotation at 40°, 1Hz. **B.** Statistics of velocity gain in response to one-way step head rotation for ±10° at 1 Hz, ±20° at 0.5 Hz, ±50° at 0.2 Hz, and ±100° at 0.1 Hz for stage 49-50 (N=7) and 55-57 (N=8). Data are presented as box plots (mean ± SEM) showing eye velocity gain for each stimulus condition (at 0.1Hz: st49-50 = 0.032 ± 0.007 *vs* st55-57 = 0.082 ± 0.015, *p* = 0.014; at 0.2Hz: st49-50 = 0.07 ± 0.015 *vs* st55-57 = 0.12 ± 0.013, *p* = 0.035; at 0.5Hz: st49-50 = 0.12 ± 0.03 *vs* st55-57 = 0.35 ± 0.04, *p* = 0.0023; at 1Hz: st49-50 = 0.21 ± 0.05 *vs* st55-57 = 0.45 ± 0.05, *p* = 0.0082; Mann-Whitney). C. Linear ratio between left and right eye velocity in response to one-way step head rotation at stage 49-50 (light brown) and 55-57 (dark brown) and corresponding linear regression lines. **D.** Example of angular VOR-driven eye position (pos.) and velocity (vel.) traces (D_1_) and averaged eye velocity (D_2_; ±SD; 20 cycles averaged) evoked by horizontal sinusoidal head rotations (head position in dark plain line, velocity in grey dashed line) at 0.5 Hz, 40° (±20°) in a stage 49-50 (light brown) and 55-57 (dark brown) larvae. **E.** Statistic of eye velocity during sinusoidal head rotations for ±10° at 1 Hz, ±20° at 0.5 Hz, ±50° at 0.2 Hz, and ±100° at 0.1 Hz. Data are presented as box plots (mean ± SEM) showing eye velocity for each stimulus condition (at 0.1Hz: st49-50 = 3.064± 0.47 vs st55-57 = 3.575 ± 0.409, p = 0.4698; at 0.2Hz: st49-50 = 4.639 ± 0.808 vs st55-57 = 6.788 ± 0.912, p = 0.0939; at 0.5Hz: st49-50 = 6.146 ± 0.938 vs st55-57 = 10.0 ± 0.794, p = 0.0111; at 1Hz: st49-50 = 5.467 ± 1.086 vs st55-57 = 10.32 ± 1.193, p = 0.0229; Mann-Whitney). **F.** Linear ratio between left and right eye velocity in response to sinusoidal head rotation at stage 49-50 (light brown) and 55-57 (dark brown) and corresponding linear regression lines. **G.** Single cycle (G_1_) and multiple cycle (G_2_, 20 consecutive cycles) raw angular eye position and associated polynomial fit of VOR-driven eye movement during sinusoidal rotation. In G_2_ light and dark brown traces represent cycles from a stage 49-50 and 55-57, respectively. **H.** Mean coefficient of determination (R^2^) calculated form to polynomial fit in G2 for each animal and represented as box plots for each stimulation frequency at stages 49-50 and 55-57. **I.** Mean phase relationship between head and eye velocity (mean ± 95% confidence interval) at stages 49-50 and 55-57 across stimulation frequencies at 60°/s.

### Central origin of the angular vestibulo-ocular maturation

The origin for the functional maturation of the angular VOR could be either peripheral or central. Indeed, the continuous growth of horizontal semi-circular canals could induce a better canal sensitivity and consequently an improved VOR response at stage 55-57 than at stage 49-50. This mechanism already determines the onset of the angular VOR (Lambert et al. 2008). However, neuronal changes in vestibulo-ocular circuits could also contributed to the VOR maturation, independently of the canal growth. First, the size of the horizontal semi-circular canals was measured from the onset of the angular VOR, at stage 49-50 to late larval stages 58. Despite a small significant increase between stage 49-50 and stage 52, horizontal canal circuit and lumen radii were comparable from stage 52 to stages 58 (Fig. 2A; Schneider-Soupiadis et al. 2025), suggesting that the canal size parameter might not be the principal reason for the VOR maturation during this developmental period. To test the hypothesis of a central origin for the angular VOR maturation, galvanic vestibular stimulation (GVS) of the horizontal canal vestibular nerve branch (Gensberger et al. 2016) was performed to induce VOR-driven eye movements, independently of the canal size (Fig. 2B-D). GVS evoked stronger and more reliable ocular responses at later developmental stages than at early stages, when the VOR started (Fig. 2B-C). Eye angular amplitude was significantly larger at stage 55-57 than at stage 49-50 for stimulations at 1Hz, 250µA (4.91°±0.9362 at stage 49-50 and 13.08°±1.48 at stage 55-57; *p* = 0.0034) and 300µA (7.947°±1.382 at stage 49-50 and 14.62°±1.381 at stage 55-57; *p* = 0.0109) and at 0.5Hz 300µA (Fig. 2D; Paired t-test). However, low amplitude GVS-evoked ocular responses were comparable between early and late larval stage (Fig. 2D), suggesting that the VOR maturation could be less important for lower stimulus frequency, as shown in figure 1B, E. Nonetheless, these results indicated clearly that the improvement of VOR was supported, at least partly, by a maturation in vestibulo-ocular circuits, potentially at the level of extraocular MN that activate horizontal eye muscles. Consequently, we investigated evidences for a potential functional maturation in abducens motoneurons between stage 49-50 and stages 55-58. First to consider, neurogenesis could be at the origin of this maturation by adding new Abducens motoneurons from the existing pool at stage 49-50. To test this possibility, a BrdU pulse-chase was performed from stage 49-50 to stage 55-57. Stage49-50 tadpoles were incubated in 10µM BrdU during 24h then were chased until stages 55-57, where Abducens motoneurons were retrogradely labelled from the lateral rectus, combined with the BrdU immunolabelling (Fig. 2E). Only a very small fraction of Abd MN was BrdU-positive (0.84% ± 0.5432, N = 7 animals), showing that almost no motoneuron innervating the muscle at late larval stage originated from mitotic cell at stage 49-50 (Fig. 2E). This indicated that neurogenesis was no longer ongoing in the abducens motor nucleus during that period and did not participate, at least significantly, in the VOR maturation. Concomitantly, the abducens soma volumes were distributed similarly from stages 49-50 to stage 55-57 (24.29% at stage 49-50, n=6, and 21.79% at stage 55-57, n=6, respectively; Fig. 5F), with a mean volume peak around ∼250 µm³. Moreover, the total number of retrogradely labelled abducens motoneurons was comparable between the two larval stages (*stages 49-50*: 48.667 ± 4.435; *stages 55-57*: 51.267± 2.363; inset inn Fig; 2F), indicating that motoneuron number and soma size did not changed during this period. Altogether, these results revealed that the maturation of the angular VOR do not rely on neurogenesis activity but would be accompanied by post-differentiation functional changes in vestibulo-ocular circuits, probably at the level of extraocular motor plant.

**Figure 2.**
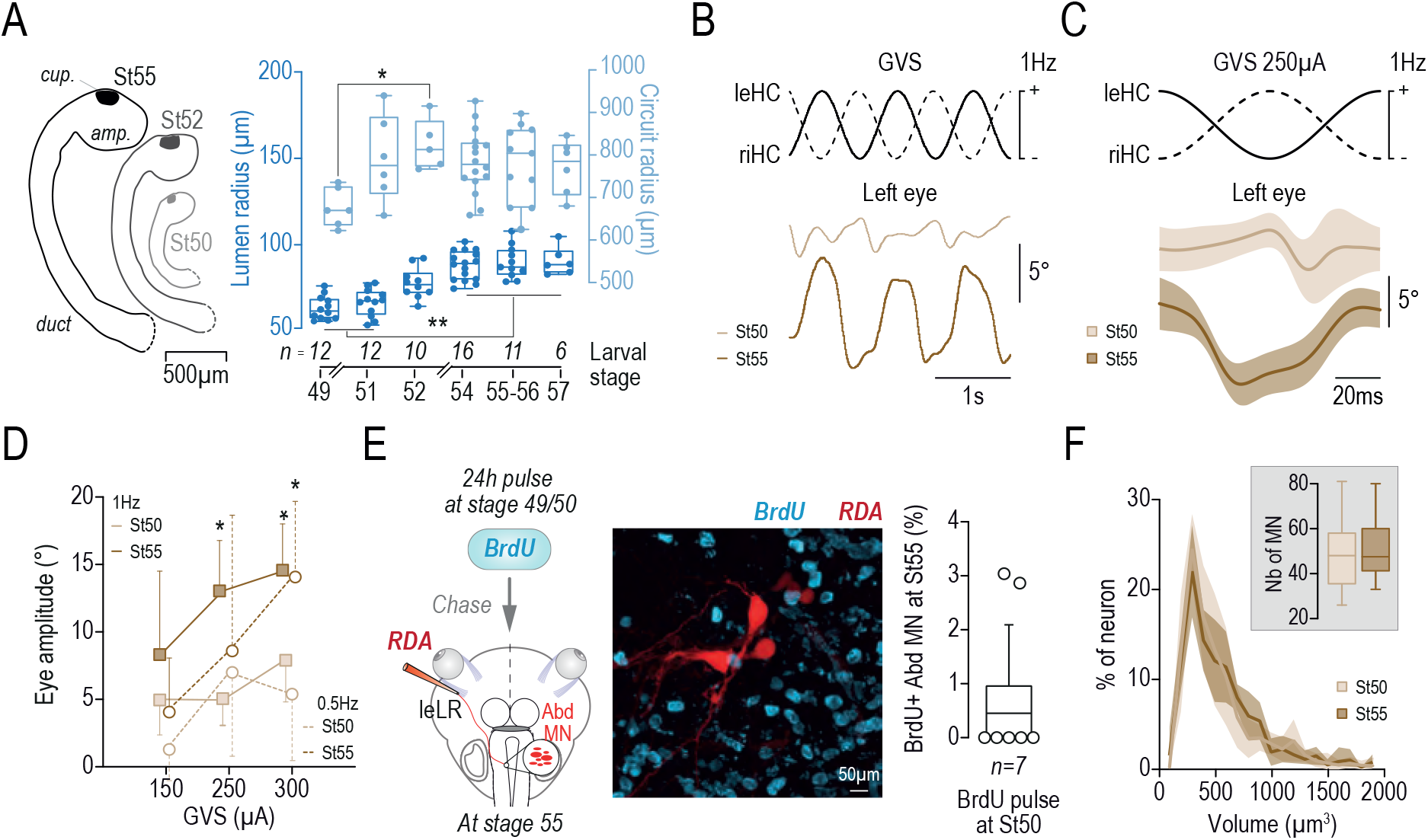
VOR maturation do not depend of canal growth and motoneuron neurogenesis. **A.** Outline of the horizontal semicircular canal (HCs) at stages (St) 50, 52, and 55 and average of horizontal canal lumen and circuit radius across developmental stages after the onset of the aVOR (mean ± SD, Kruskal Wallis. Circuit radius: St49 vs 52, p= 0.0412. Lumen radius: St 49 vs St54, 56, 57-58, *p* <0.0009; St 51 vs St54, 56, 57-58, *p* < 0.0046). **B.** Representative example of horizontal eye movement evoked by galvanic vestibular stimulation (GVS) of horizontal canal cupula (250µA, 1Hz) at stages 49-50 and 55-57. **C.** Average (±SD; over 20 cycles) cyclic ocular response evoked by GVS at stages 49-50 and 55-57. **D.** Eye amplitude (°; mean ± SEM) evoked by GVS at 0.5 and 1 Hz and 150, 250, and 300 µA (stage 49-50: n = 5; stage 55-57: n = 7; p values =0.0034 and 0.0109 respectively). **E.** Schematic view of the BrdU pulse-chase experiment (left side), confocal image of Abducens MN labelled with rhodamine dextran amine (RDA) and immunolabeling of BrdU+ cells; proportion of Abducens MN at stage 55 deriving from a mitotic cell at stage 49-50 (0.84% ± 0.5432; n = 7). **F.** Distribution of Abducens MN soma volume at stages 49-50 and 55-57 (n=6 at both stages; no significant differences from stage 49-50 and stage 55; Kolmogorov-Smirnov). Top right inset: total number of retrogradely labelled Abd MN counted on transverse sections at stages 49-50 and 55-57 (48.667 ± 4.435 and 51.267± 2.363; n =15 and 30 animals per stage).

### Discharge components in vestibular-induced abducens motor response

To investigate the possibility of functional changes at the extraocular motor plant related to the maturation of angular VOR, the discharge of the abducens motor nerve branch innervating specifically the lateral rectus muscle (Fig. 3A) was analysed at stages 49-50 and 55-57 in response to head rotations. Abducens motor units were previously characterized in larval frog according to their activity at rest (Dietrich et al. 2017). Spontaneous single units exhibited a resting discharge with a firing rate above 1 spikes/s whereas silent units did not fire at rest or with a rate below 0.1 spikes/s for most of them (Fig. 3A). Supposedly, spontaneous and silent abducens units innervated multiple innervated and single innervated muscle fibres, respectively (Fig. 3A, (Dietrich et al. 2017). Also, silent motor unit exhibited spikes with a higher amplitude than spontaneous motor units (Fig. 3B; (Dietrich et al. 2017). Using the same criteria than Dietrich and colleagues (2017), we identified two multi-unit components in the abducens nerve discharge, one spontaneously active at rest (blue area in Fig. 3B) and one almost exclusively active during head movement only (orange area in Fig. 3B). Spikes of the spontaneous multi-unit component fired at rest with a firing rate of 91.1±12.6 spikes/s (Fig. 3C) and a spike interval distributed from a few milliseconds to 400ms (Fig. 3D). Inversely, spikes for the silent component of the nerve discharge fired at rest with a rate 100 time lower than spontaneous one (1.015±0.336 spikes/s; Fig. 3C) and a spike interval comprised between 800ms and 37600ms (Fig. 3D), demonstrating that abducens motor units of this component were silent at rest. Like the activity of the silent motor units was motion-dependant; the dynamic of both silent and spontaneous components of the abducens discharge were analysed first in response to either yaw (Fig. 3E) or roll (Fig. 3F) axis head rotations. At stage 55, the firing rate of the spontaneous discharge component increased during a horizontal (yaw) one-way step of the head with a peak discharge occurring around the end of the displacement when the head reached its final position (65.7 ± 33.0ms; Fig. 3E_1-2_, left plot). After the peak, the firing rate of the spontaneous component decreased slowly with a half-response-decrease latency of 376.1 ± 61.68ms (Fig. 3E_1-2_, right plot). During the same head motion stimulus, the firing rate of the silent motor unit group increased rapidly with a peak response occurring at 178.7 ± 17.9ms before the end of the step displacement (Fig. 3E_1-2_, left plot); much faster than the spontaneous component. Also, the half-response-decrease latency was shorter for the silent than for the spontaneous discharge component (237.1 ± 37.68ms; Fig. 3E_1-2_, right plot). The spontaneous unit group presented the same low discharge dynamic during horizontal step at stage 49-50 and stage 55-57 (Fig. 3E_3_), with a broad phase relationship non-specifically locked on either the head position or velocity signal (Fig. 3E_4_). Inversely, the silent discharge component improved during the development demonstrating a faster and sharper response dynamic, better phase locked on the head velocity signal during horizontal rotations at stage 55-57 compared to stage 49-50 (Fig. 3E_4_).

**Figure 3.**
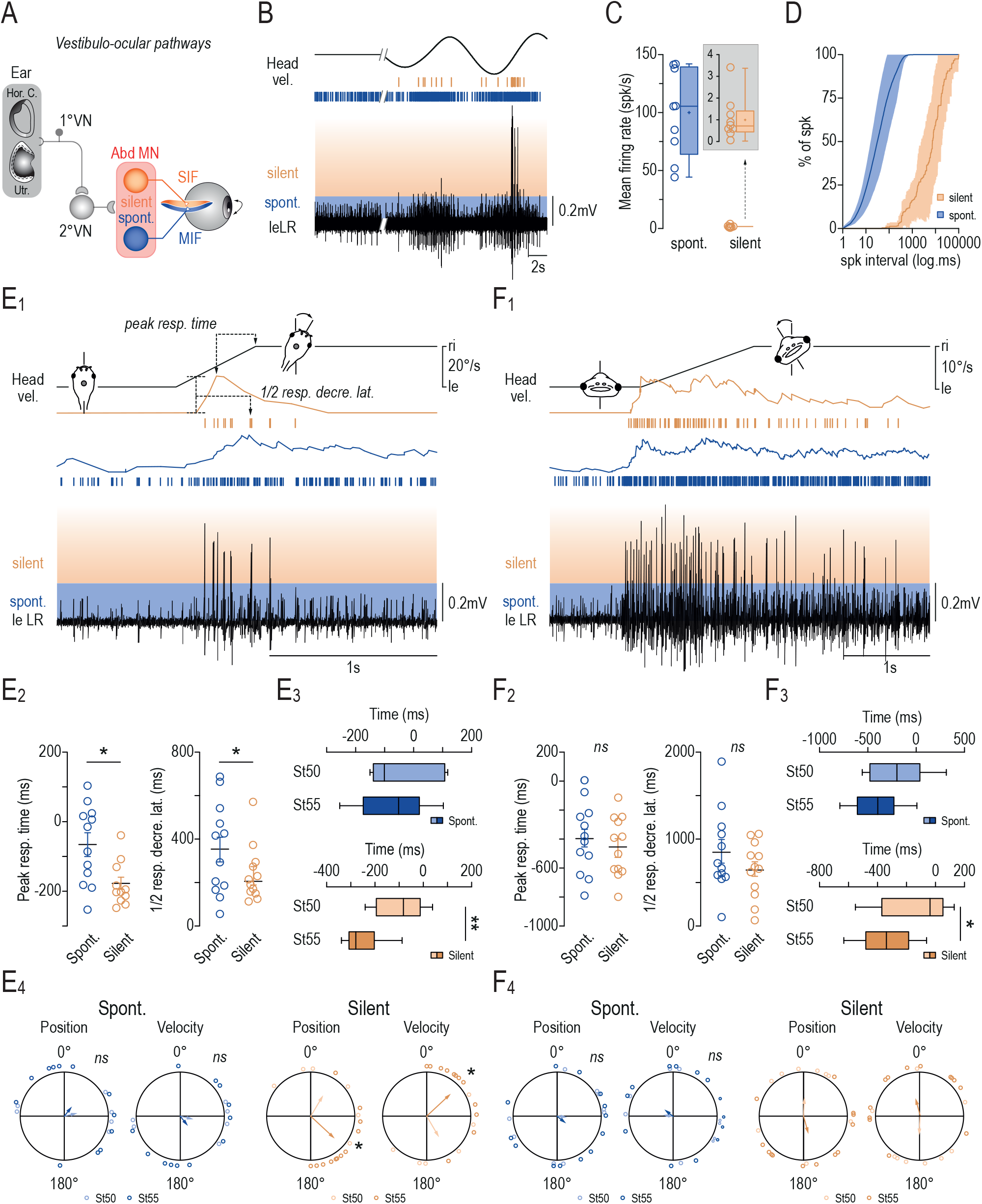
Vestibular-induced responses dynamics of abducens multi-unit components. **A.** Simplified schematic view adapted form Dietrich et al., (2017) of VOR sensory-motor pathway contacting silent and spontaneous (spont.) Abducens motoneurons subpopulations that innervate singly innervated fibers (SIF) and multiply innervated fibers (MIF), respectively. **B.** Extracellular recording of the abducens motor nerve activity at rest and during horizontal head rotation in a stage 55 tadpole. Blue and orange bands delimit the spontaneous (blue) and silent (orange) multi-unit components of the Abducens discharge. Vertical blue and orange events above the nerve trace indicate all spikes detected for each discharge component. **C.** Box plot firing rate in spike/s (spk/s) of spontaneous and silent multi-unit discharge components at rest (n=9). The firing rate range of silent component is indicated in the inset. Upper and lower error bars represent maximum and minimum values, respectively. Bounds of box and center lines represent 25, 75 % percentile and median values, respectively. Superimposed full colored circles represent mean firing rates. **D.** Cumulative distribution (±SD) of spike intervals in spontaneous and silent component at rest. **E-F.** Response dynamics of spontaneous and silent multi-unit discharge components during one-way step head horizontal (yaw) rotation at 1Hz, 20°/s (E) and one-way step head vertical (roll) rotation at 1Hz, 10°/s (F). Means (±SEM) of peak response time and half response decrease latency (1/2 resp. decre. Lat.) of spontaneous and silent discharge components during yaw and roll head turns are represented in E_2_ and F_2_, respectively. Box plots (same parameters than in C) of peak response time and half response decrease latency from both spontaneous and silent components during yaw and roll head turns at stage 49-50 (light blue n=6, light orange; n=10) and 55-57 (blue and orange; n=12) are represented in E_3_ and F_3_, respectively. The phase relationships of the peak response for both spontaneous and silent components and for both stage 49-50 and 55-57 with either the position or the velocity peak of the head turn are indicated with polar plots in E_4_ (yaw) F_4_ (roll), respectively. *p = 0.0015,* Wilcoxon in E_2_ left; *p = 0.021,* Mann-Whitney test in E_2_ right; *p = 0.0047,* Mann-Whitney in E_3_ bottom; *p = 0.0249,* Mann-Whitney in F_3_ bottom*; p = 0.002,* Watson-Williams test for both plots in E4. Hor. C. = horizontal canal; St = stage; Utr. = utricles; Vel. = velocity; leLR = left lateral rectus muscle; 1°VN, 2°VN = 1rst and 2^nd^ order vestibular neurons.

Spontaneous units responded broadly to one-way step of the head in roll axis at both stage 49-50 and 55-57 (Fig. 3F), with a peak response time around 400ms (396.8 ±70.34ms at stage 55) before the end of the step displacement. The half-response-decrease latency was quite long for this abducens discharge component (847.2 ± 136.6ms; Fig. 3F_2-3_). As long as the head was maintained in the tilted position, the discharge rate of the spontaneous unit group remained higher than the resting rate before the tilt, demonstrating a constant activation by utricle inputs (Fig. 3F_1_). The spontaneous discharge peak was not specifically phase-locked on neither the position nor the velocity signal of the head motion (Fig. 3F_4_). Silent units also responded broadly to head roll tilt with a peak response occurring around 454.8 ± 60.39ms before the end of the step, comparably to spontaneous units (Fig. 3F_2-3_). However, the peak response occurred later in stage 49-50 (195.6 ± 88.2ms) than in stage 55-57 (Fig. 3F_3_). The half-response-decrease latency for the silent component was also quite long but still shorter than the spontaneous one and the discharge rate came back to its resting baseline after few seconds despite the maintain of the head tilted, suggesting a weaker activation of silent motor units by utricles inputs. Like the spontaneous component, the silent discharge peak was not particularly phase-locked on neither position nor velocity signal of the head motion (Fig. 3F_4_). Altogether, these results confirmed that the abducens motor nerve discharge presented two groups of units, constituting two different components in the nerve discharge. In addition, our findings revealed that these two groups of units encoded differently head motion parameters during horizontal or vertical tilts. Spontaneous units presented a tonic discharge, with a slow dynamic, related to the head position and receiving massive utricular inputs whereas silent unit fired with a fast response dynamic, related to the head velocity signal. This dual slow-tonic or fast-phasic discharge dynamic in abducens units, encoding specifically either position or velocity signals, was previously reported in adult frog in response to optokinetic step stimulations (Dieringer et Precht 1986). In addition, only the silent component demonstrated a significant improvement of its firing capacity in response to horizontal head tilts during the larval development.

### Differential changes in abducens nerve discharge dynamics during aVOR maturation

The functional maturation of both spontaneous and silent unit groups was characterized more extensively by analysing their discharge modulation in response to sinusoidal horizontal rotations of the head over the larval development (Fig. 4A_1-2_). During sinusoidal head rotations, the discharge of the abducens motor nerve increased regularly when the head moved to the opposite side (ON phase) of the recorded nerve and became silent when the head turned back to the side of the nerve (OFF phase; see example in Fig. 4A with the abducens nerve innervating the left lateral rectus). In this stimulus configuration, the firing rate of the spontaneous multi-units component increased during the ON phase the same way in both stage 49-50 and 55-57, reaching a maximum of ∼75 spikes/s and ∼60 spikes/s for low (0.2Hz; Fig. 4B_1_) and high frequency (1Hz; Fig. 4B_2_), respectively. The firing modulation of the spontaneous unit group was comprised between 60 and 40 spikes/s during increasing head rotation frequencies from 0.1 to 1Hz (at constant 30°/s head velocity; Fig. 4C_1_) and increased from 5-10 spikes/s to 100 spikes/s during increasing head velocities (at constant 1Hz frequency; Fig. 4C_2_). This spontaneous firing modulation did not show any significant variation from stage 49-50 to stage 55-57, except at 0.1Hz (Fig. 4C_1-2_). Inversely, the firing rate of the silent multi-unit component at the ON phase was much higher in old larvae than in early one, twice the maximum rate at low frequency (0.2Hz; Fig. 4D_1_) and even more at high frequency (1Hz; Fig. 4D_2_). The firing modulation for the silent unit group was constant independently of the head frequency, but significantly higher at stage 55-57 (approx. 30 spikes/s) than at stage 49-50 (approx. 10-20 spikes/s; Fig. 4E_1_). During head velocity range, the silent firing modulation increased linearly but at a significantly higher rate in old larvae than in early one, except for very low velocities (Fig. 4E_2_).

**Figure 4.**
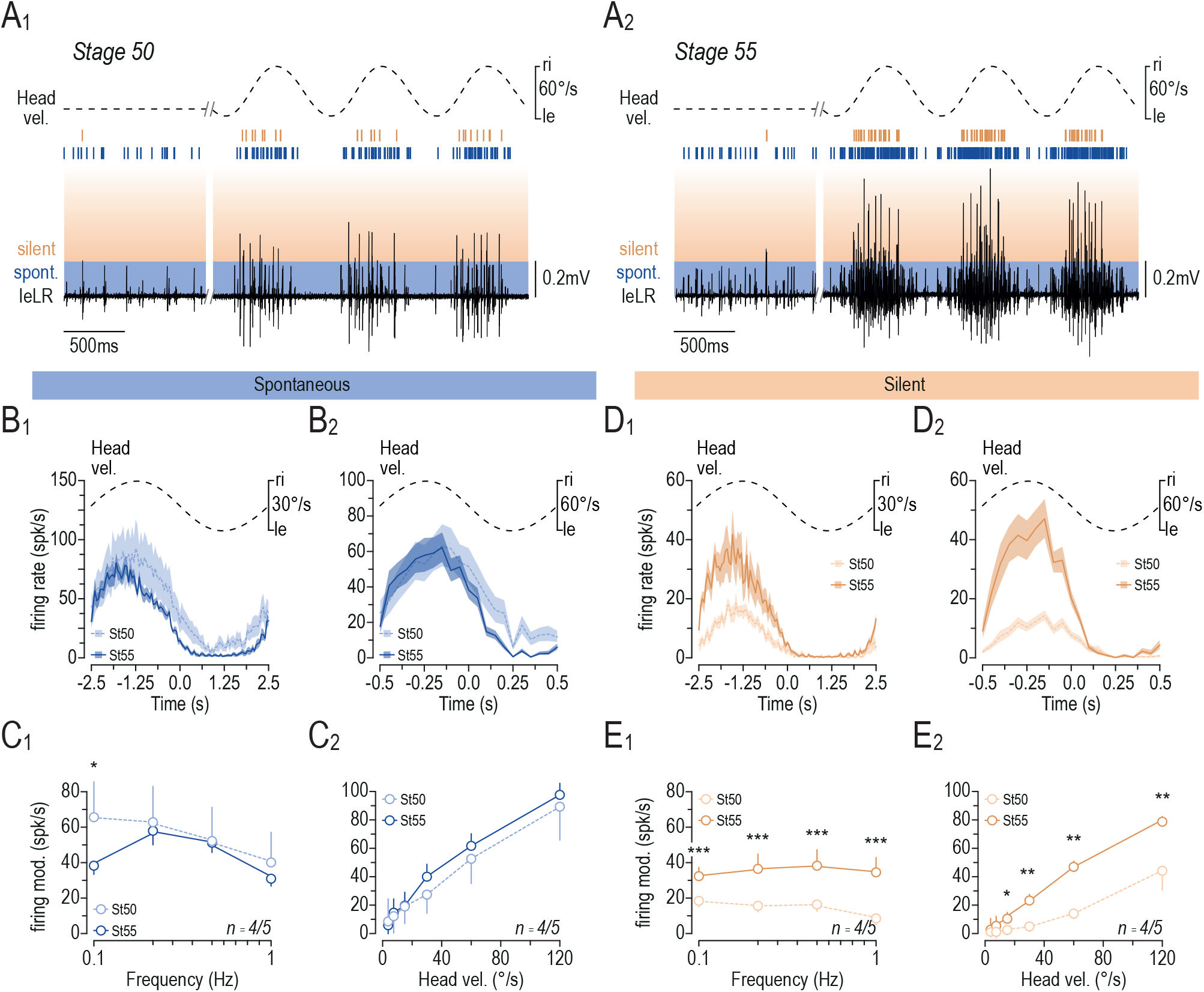
Firing modulation of spontaneous and silent discharge multi-unit components in Abducens motor activity during larval development. **A.** Extracellular recording of the abducens motor nerve activity at rest and during sinusoidal horizontal head rotation (1Hz, ±60°/s) in a stage 50 (A_1_) and a stage 55 (A_2_) tadpole. Blue and orange bands delimit the spontaneous (blue) and silent (orange) multi-unit components of the Abducens discharge. Vertical blue and orange events above the nerve trace indicate all spikes detected for each discharge component. **B.** Averaged (±SD) firing rate of spontaneous multi-unit discharge component at stage 50 (light blue) and 55 (blue) in response to sinusoidal horizontal head rotation at 0.2Hz ±30°/s (B1) and 1Hz ±30°/s (B2). **C.** Bode (C_1_, at ±30°/s) and linearity (C_2_, at 1Hz) analysis of averaged (±SEM) firing modulation (mod.) of spontaneous multi-unit discharge component at stage 50 and 55. **D.** Averaged (±SD) firing rate of silent multi-unit discharge component at stage 50 (light orange) and 55 (orange) in response to sinusoidal horizontal head rotation at 0.2Hz ±30°/s (D1) and 1Hz ±30°/s (D2). **E.** Bode (E_1_, at ±30°/s) and linearity (E_2_, at 1Hz) analysis of averaged (±SEM) firing modulation of silent multi-unit discharge component at stage 50 (n=4) and 55 (n=5). * *p<0.05, ** p<0.01, *** p<0.005* in E_1-2_, Mann-Whitney U test. LR = lateral rectus; Ri/le = right/left; St = stage; Vel. =velocity

Results shown in figure 4 were confirmed by analysing the discharge of spontaneous and silent single units identified by spike sorting screening of the multi-unit recording (Figure 5). For each preparation, several spontaneous and silent single units were clearly identified, based on the spike shape and amplitude (Fig. 5A-B). Only single units constituting a clear homogenous population sorted out from the principal component analysis (Fig. 5C) were considered for discharge analysis. Very small amplitude units presented too much variations of the spike shape due to the effect of the baseline noise and were excluded from the study. Usually, 5-12 spontaneous and 3-6 silent single units were identified from a multi-unit recording through the spike sorting analysis (see the example shown in Fig. 5A-D). The firing rate of these single units was calculated at all stimulus frequencies and velocities (see example in Fig. 5D). The ON phase firing rate of spontaneous single units was identical at stage 49-50 and stage 55-57, independently of the head frequency or velocity (Fig. 5E_1-2_). At both larval stages, the firing modulation of the spontaneous units was around 15-20 spikes/s during increasing stimulus frequencies from 0.1 to 1Hz (at 30°/s head velocity; Fig. 5F_1_) and increased from ∼5 spikes/s to ∼25 spikes/s during increasing head velocities (at 1Hz frequency; Fig. 5F_2_), comparable to results shown in Figure 4. Inversely, silent single units demonstrated a firing rate during the ON phase significantly higher at stage 55-57 than at stage 49-50 (Fig. 5G_1-2_). At stage 49-50, the firing modulation was constant around 10 spikes/s at 0.1-0.5Hz and dropped off to 5 spikes/s at 2Hz but increase from 15 spikes/s up to 20 spikes/s at stage 55-57, even at 2Hz (Fig. 5H_1_). During head velocity range, the firing modulation of silent single units increased linearly but also at a higher rate in late larvae than in early one, except for very low velocities (Fig. 5H_2_), also confirming the results shown in Figure 4. Altogether, results obtained from the multi-unit and single unit analysis of the lateral rectus motor nerve discharge revealed abducens motoneuron sets showed different maturation profiles during larval life (Figures 4 and 5). From stage 49-50, only silent abducens motor units exhibited a significant increase of their discharge modulation in response to head rotations, demonstrating that this set of abducens motoneurons become more responsive to head movements detection by semi-circular during this late larval period. Inversely, spontaneous motor unit were modulated with the same amplitude at both early and late larval stages, suggesting that maturation no longer occurs in this motoneuron subgroup during this developmental period but might occurred earlier.

**Figure 5.**
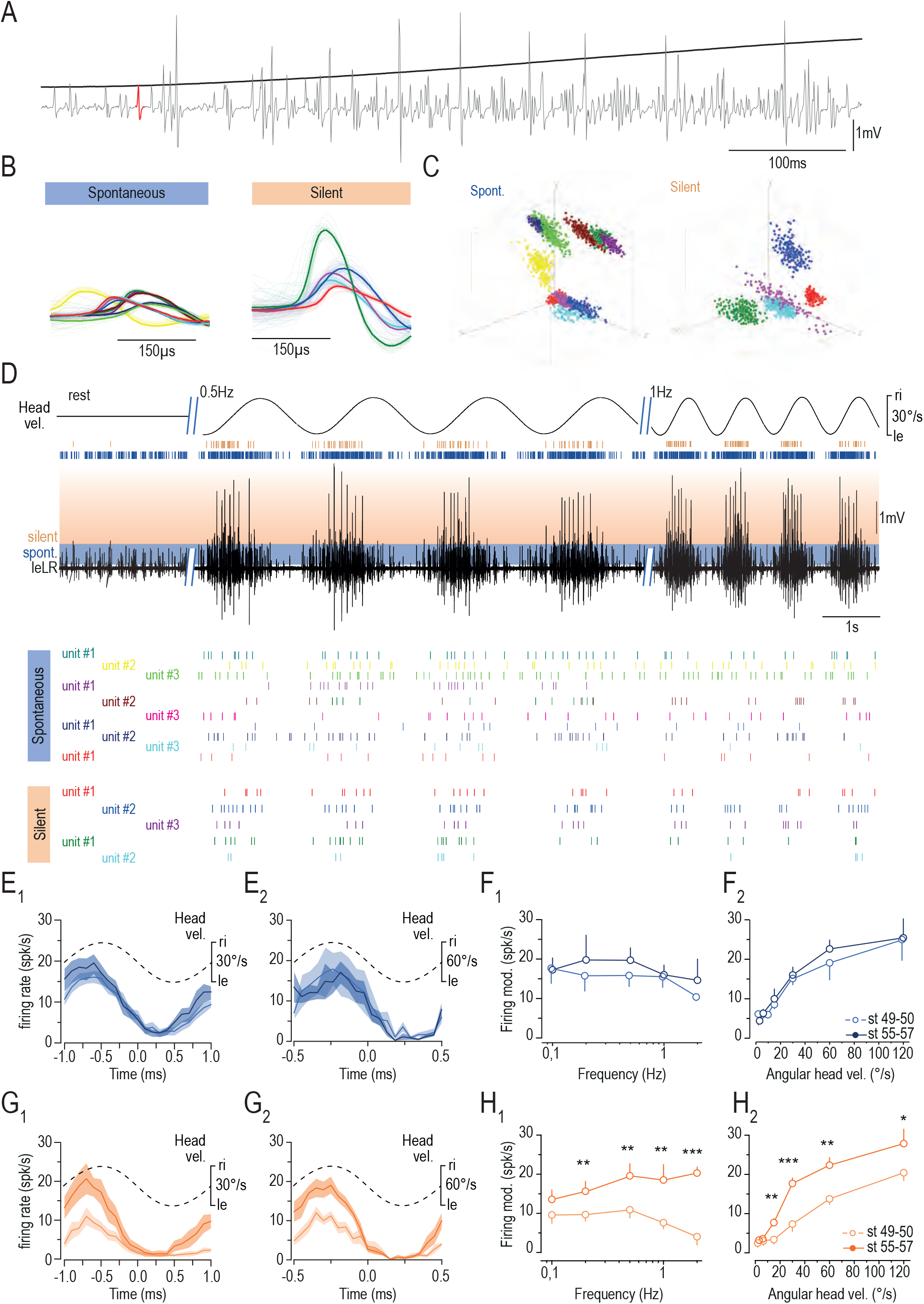
Firing modulation of spontaneous and silent Abducens motor single units during larval development. **A.** Detail of the extracellular recording of the Abducens motor nerve during horizontal head rotation at stage 55 showing the diversity of motor units composing the nerve activity. **B-C.** Shape spike templates (B, the thick line is the average of thin lines) and corresponding clusters identified with principal component analysis (C) of spontaneous and silent single units extracted from the recording showed above. **D.** Extracellular recording of the abducens motor nerve activity at rest and during sinusoidal horizontal head rotation (0.5Hz and 1Hz ±30°/s) in a stage 55 tadpole. Blue and orange bands delimit the spontaneous (blue) and silent (orange) multi-unit components of the Abducens discharge. Vertical blue and orange events above the nerve trace indicate all spikes detected for each discharge component. Coloured vertical events below the trace indicate all spikes detected from the different single units identified in B-C. **E.** Averaged (±SD) firing rate of spontaneous single units at stage 50 (light blue) and 55 (blue) in response to sinusoidal horizontal head rotation at 0.5Hz ±30°/s (E_1_ left) and 1Hz ±60°/s (E_2_ right). **F.** Bode (F_1_, at ±30°/s) and linearity (F_2_, at 1Hz) analysis of averaged (±SEM) firing modulation (mod) of spontaneous single units at stage 50 and 55. **G.** Averaged (±SD) firing rate of spontaneous single units at stage 50 (light blue) and 55 (blue) in response to sinusoidal horizontal head rotation at 0.5Hz ±30°/s (G_1_ left) and 1Hz ±60°/s (G_2_ right). **H.** Bode (H_1_, at ±30°/s) and linearity (H_2_, at 1Hz) analysis of averaged (±SEM) firing modulation of spontaneous single units at stage 50 (n=4) and 55 (n=5). * *p<0.05, ** p<0.01, *** p<0.005* in E_1-2_, Mann-Whitney U test. LR = lateral rectus; Ri/le = right/left; St = stage; Vel. =velocity.

### Structural changes of the abducens nerve and of the lateral rectus innervation during angular VOR maturation

Electronic microscopy was performed on abducens motor nerve ultra-thin cross sections (Fig. 6A-B) to determine if the functional maturation in motor units was associated with structural changes between early and late larval stages. At both stages, the nerve contained myelinated axons identified by their dark myelin sheaths (see insets “a” in Fig. 6A-B) as well as unmyelinated axons (see insets b in Fig. 6A-B). The number of myelinated axons increased significantly between stages 49-50 and 55-57 (from 2.8±1.393 to 19.63±1.880 axons; Mann-Whitney, p = 0.0008; Fig. 6C, left plot). Additionally, the myelinated axonal surface (from 0.9±0.47 to 8.92±1.38 µm^2^ axons surface; Mann-Whitney, p = 0.0010; Fig. 6C, middle plot) and the surface of myelin sheath (from 0.378±0.18 to 2.98 ±0.468 µm^2^ myelin surface; t-test, p =0.0016; Fig. 6C, right plot) were also significantly larger at later stages. Inversely, the number of unmyelinated axons remained unchanged (Fig. 6D, left plot). However, their axonal surface increased modestly (from 0.232±0.018 to 0.365±0.054 μm^2^, p = 0.0420; Mann-Whitney; Fig. 6D, right plot) but in a less extent than myelinated axons. These results demonstrated an important improvement of fast-conducting axonal fibres in the abducens motor nerve, likely corresponding to silent motor units.

**Figure 6.**
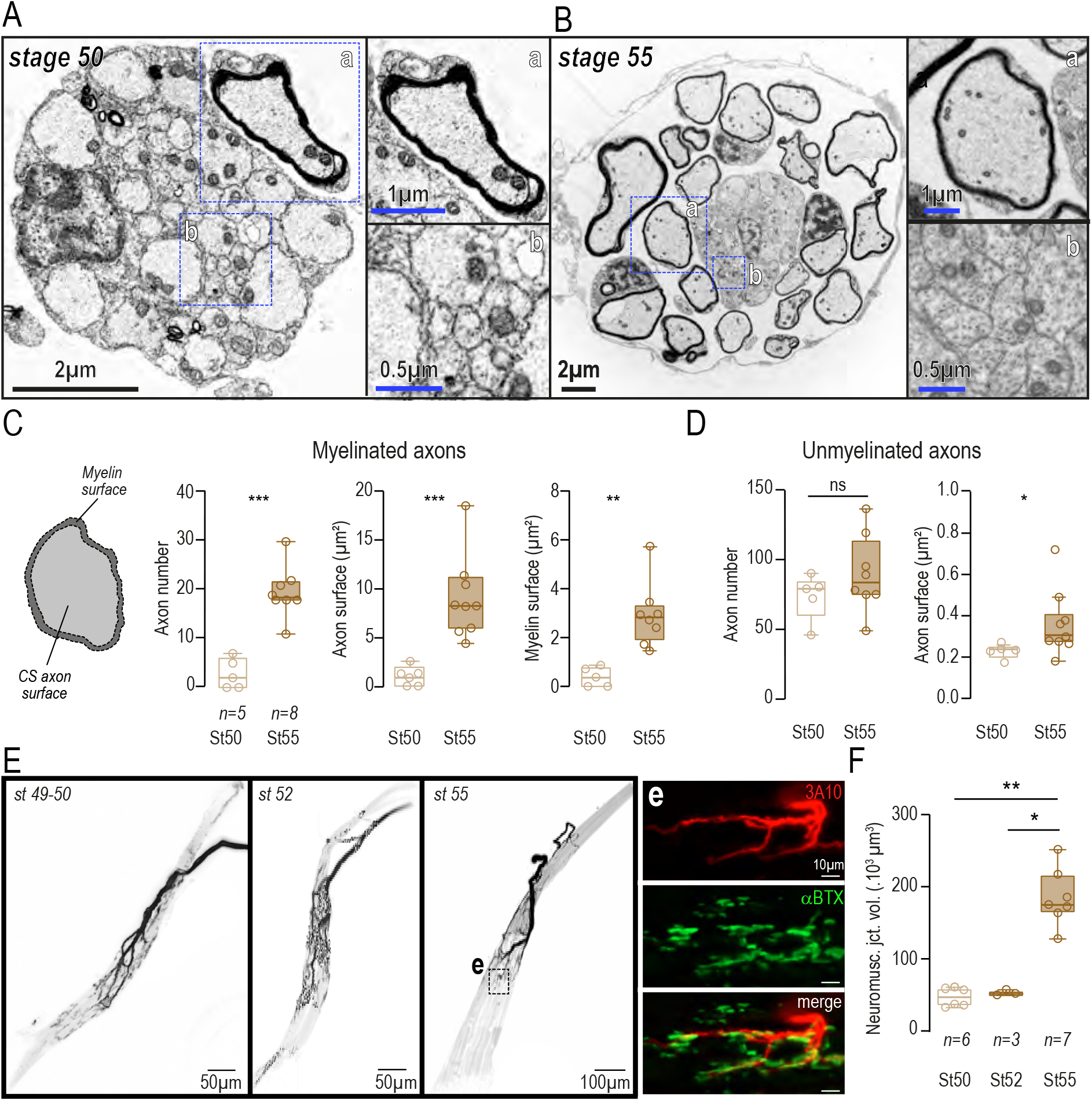
Axonal structure of the Abducens motor nerve during the larval development. **A-B.** Electronic microscopy images of ultra-thin coronal sections of an Abducens motor nerve at stage 50 (A) and stage 55 (B). Inset pictures a and b show a magnified view of myelinated (a) and unmyelinated (b) axons, respectively. **C-D.** Box plots of axon number, axon surface and myelin surface for myelinated (C) and unmyelinated (D) axons at stage 50 (N=5) and stage 55 (N=8). The left scheme in C indicates how was calculated the surfaces. Upper and lower error bars represent maximum and minimum values, respectively. Bounds of box and center lines represent 25, 75 % percentile and median values, respectively. In C: p=0.0008 (left), p= 0.0010 (middle), p=0.0016 (right), Mann-Whitney; in D: p= 0.0420, Mann-Whitney. **E.** Confocal stack images of a lateral rectus muscle with the Abducens motor nerve insertion at stage (St) 49-50, 52, and 55. The inset picture “e” in the right image shows a x40 magnified view of the neuromuscular junction plates on the muscle. The nerve is labelled with an immunostaining against neural tubulin (3A10) and the neuromuscular junction with the α-bungarotoxin (αBTX). **F.** Box plots of confocal stack volume occupied by neuromuscular junctions (Neuromusc. Jct. vol) on the lateral rectus muscle at stage 50 (N=6), 52 (N=3), 55 (N=7). p=0.0012 for 50vs55 and p= 0.0167 for 52vs55; Mann-Whitney.

Such a structural improvement would conduct to a faster transmission of the output signal in this subset of motoneurons, corroborating the functional maturation of silent motor units from early to late larval stages. Concomitantly, the absence of change in unmyelinated fibres, characterized by a slow conducting signal transmission, could be related to the absence of functional maturation observed in spontaneous motor units during the same period. These axonal modifications were accompanied with an improvement of the innervation of the lateral rectus muscle by the Abducens motor nerve quantified by the volume of nerve-muscle contact (Fig. 6E). Combined neurofilament immunolabeling and α-bungarotoxin labelling allowed to visualize the total amount of nerve-muscle contact on whole-mount lateral rectus muscle and Abducens nerve. No difference of contact volume was observed between stages 49-50 and 52 (46.7×10³±5.3 μm³ and 52.8×10³±2.2 μm³ respectively; N=6 and N=3, Mann-Whitney, p>0.05). However, a significant increase in nerve-muscle contact was detected at stage 55-57 (184.5×10³±14.9 μm³, n = 7, Mann-Whitney, p =0.0012 vs stage 49-50 and p=0.0167 vs stage 52; see Figures 6E-F), demonstrating a better innervation of the muscle by the abducens nerve between stage 52 and 55-57. Altogether, these results revealed that the functional maturation observed for silent motor units was accompanied by anatomical changes in the motor nerve enhancing a better output signal transmission and a better muscle activation.

## DISCUSSION

In xenopus tadpoles, two functional types of Abducens motoneurons participate to the motor command driving conjugate compensatory eye movements that constitutes the angular VOR: motor units spontaneously active at rest and motor units silent at rest. During the larval period, the angular VOR matured until the end of the pro-metamorphosis by providing a better control of the eye velocity. This improvement of the ocular kinematic in response to head rotations was sustained by a better firing modulation of silent Abducens motoneurons only, while the firing modulation of spontaneous motor units remained unchanged. This functional maturation in silent Abducens motoneurons was accompanied by an increase of the myelination in the abducens motor nerve and a larger innervation of the lateral rectus muscle, allowing a faster signal conductivity and a stronger muscular activation for a same sensory input.

### Conjoint maturation of silent abducens motoneurons with the angular VOR

In larval amphibians, the maturation of the angular VOR was briefly reported in previous publications by an increase of the reflex gain (Lambert et al. 2008; Bacqué-Cazenave et al. 2022; Schneider-Soupiadis et al. 2025). Comparable maturation pattern was also described in fish either in response to vertical or horizontal head rotations, in early or late larval life, respectively However, this developmental mechanism remained poorly investigated. The present study characterized more precisely the behavioral changes in the VOR-driven ocular kinematic to demonstrate that the VOR maturation conducted to a better control of the eye velocity. The canal duct morphology is a critical parameter in the vestibular sensitivity dedicated to the detection of the head rotation and consequently to the developmental onset of the angular VOR in aquatic species (Lambert et al. 2008; Beck et al. 2004; Schneider-Soupiadis et al. 2025; Lambert et Bacqué-Cazenave 2020). Nevertheless, it cannot be excluded that changes in central vestibulo-ocular pathways also participate in the maturation of the angular VOR. Recent works in early larval zebrafish, support this idea for the development of the otolith-driven VOR (Goldblatt et al. 2023, 2024; Leary et al. 2025). Therefore, it is likely that comparable developmental mechanisms are involved in the maturation of the angular vestibulo-ocular reflex (VOR) in larval amphibians. Indeed, our results revealed that such a central maturation occurs at least in the extraocular motoneurons. In addition, the sub-population of abducens motor neurons, silent at rest, that matured conjointly with the angular VOR, is precisely the subtype encoding for the velocity signal, the ocular kinematic parameter directly impacted by the VOR maturation. Using the same electrophysiological approach, a past preliminary study reported comparable adaptation of the discharge modulation in Trochlear motor units during larval Xenopus development (Westhoff 2006). Additionally, to the increase in the discharge modulation, the increase of the number of myelinated axons, combined with a thicker myelin sheath, supported the idea that only a subset of abducens motoneurons acquired their myelin sheath after stage 49-50, once the angular VOR appeared. By conducting the motor signal faster, the myelination of the abducens nerve would participate to ameliorate the eye velocity of the angular VOR-driven ocular displacements in response to the same head rotation amplitude. Furthermore, previous anatomical investigation demonstrated that the dendritic arborization and soma shape of *Xenopus* abducens motoneurons was not achieved at stage 49-50 but complete only at stage 54 (Matesz 1990).

The larger dendritic arborization should allow abducens motoneurons to receive more inputs, particularly from semicircular canal-related premotor pathways. This would consequently increase their sensitivity by raising their firing rate in response to sensory signals. Interestingly, electrophysiological and morphological changes were also reported in oculomotor motoneurons (Carrascal et al. 2006, 2009) in a temporal sequence fitting with the maturation of OKR and VOR during the rodent postnatal development (for review see Beraneck et al. 2014). Comparatively, other developmental changes could affect electrophysiological properties of abducens sub-populations in larval xenopus, like ion channel expression, modifying the motoneuron excitability and remodeling the discharge dynamic and the firing rate during head rotations. Altogether, past studies and our present work suggests the existence of a common basic developmental feature in vertebrates where extraocular motoneurons would mature conjointly to gaze-stabilizing reflexes.

### Developmental relationship in the ontogeny of extraocular motor populations and gaze-stabilizing reflexes

Whereas OKR and otolith VOR are slow dynamic reflexes producing slow eye movements for gaze-holding, angular VOR, saccades and locomotor-induced gaze response require fast ocular shifting. To insure such a diversity of gaze-stabilizing ocular behaviors, premotor circuits, treating different sort of input signals, need to project to distinct sets of extraocular motoneurons expressing appropriate morphological and physiological properties (Horn et Straka 2021). Some literature supports a model model within extraocular motor neurons are organized in functional subgroups that activate different muscle fibers subtypes to produce specific ocular kinematics. In amphibians, at least two functional subtypes of abducens motoneurons participate to the ocular kinematic produced by the angular VOR but also during the OKR (Dieringer et Precht 1986; Dietrich et al. 2017). Spontaneous abducens motor units exhibit a tonic discharge with slow dynamics, responsible for progressive angular positional adjustments of the eye. Inversely, silent motor units demonstrate a fast-phasic discharge dynamic, used to encode velocity signal. Concomitantly, these two types of extraocular motoneurons would innervate distinct types of lateral rectus muscular fibers: 1/ silent motor units connecting twitch or single innervated fibers (SIF) strictly activated during fast ocular movements and 2/ spontaneous units projecting to slow-tonic or multiple-innervated fibers (MIF; Dieringer et Precht 1986). Such a relationship between motoneurons and muscle fibers was also described in oculomotor motoneurons innervating the medial rectus muscle, the synergist muscle of the lateral rectus for horizontal ocular movements (Büttner-Ennever et al. 2001; Davis-López De Carrizosa et al. 2011; Carrero-Rojas et al. 2021; for review see Bellegarda et al. 2025). Our results confirmed this functional organization but, more interestingly, revealed that these two types of abducens motoneurons receive differently vestibular sensory inputs from otolith and semicircular canals. Whereas spontaneous motor unit are strongly activated by both canal and utricle inputs, it appeared that silent units receive limited inputs from utricles (see Fig. 3). This supports the idea that extraocular motoneurons sub-populations are activated by distinct sensory-motor vestibulo-ocular pathways (Straka et al. 2009; Horn et Straka 2021) and consequently are not involved equally in all gaze-stabilizing reflexes. Indeed, this was recently demonstrated in oculomotor motoneurons responsible for saccades either involved in visual exploration or required for prey hunting in larval zebrafish (Dowell et al. 2025). In larval frog and potentially in other animal groups, gaze motor control modalities appear and mature differently during the development according to the sensory or intrinsic signal used. Whereas OKR and otolith VOR are set early and mature rapidly, angular VOR become active in late larvae (Lambert et al. 2008; Bacqué-Cazenave et al. 2022). The LGR onset also appears early but matures conjointly with the angular VOR (Bacqué-Cazenave et al. 2022). These different maturation temporal patterns should be reflected in the extraocular motoneuron diversity, not only in the abducens nucleus but also in trochlear and oculomotor. Consequently, tonic motoneurons, with a slow dynamic discharge, should exhibit a rapid maturation from the onset of the OKR and the otolith VOR that require slow ocular kinematics. Inversely, phasic motoneurons, with a fast-dynamic discharge, seems to present a limited development until the onset of the angular VOR where they start to mature more efficiently. This would explain, at least partly, why the LGR, appearing with the OKR and otolith VOR at stage 42, exhibits a second step of maturation coordinated with the angular VOR development, conducting to a spino-ocular coupling more efficient in old larvae (Bacqué-Cazenave et al. 2022).

In conclusion, this study brought new evidences in the understanding of the functional organization of Abducens motoneurons groups and their maturation across the larval development in an animal model where basic VOR circuits are highly conserved compare to other vertebrate’s lineages. A broader investigation, combining the same methods with intracellular recordings will provide a more complete picture of the developmental pattern of extraocular motoneurons groups, a topic that remains largely unexplored today.

## Acknowledgements

This work was supported by the Centre National de la Recherche Scientifique, and granted from the Agence Nationale pour la Recherche (*ANR-22-CE16-0004-02 MOTOC* and *ANR-24-CE92-0005-01 OVOR* F.M. Lambert). Authors are grateful to Lionel Parra-Iglesias (INCIA) for taking care of the animals.

